# Dynamic hyperplastic cardiac growth in Burmese pythons

**DOI:** 10.1101/2025.05.19.654898

**Authors:** Yuxiao Tan, Thomas G. Martin, Angela K. Peter, Christopher D. Ozeroff, Christopher C. Ebmeier, Ryan Doptis, Brooke Harrison, Leslie A. Leinwand

## Abstract

Cardiomyocytes hyperplasia is the primary form of fetal heart growth, whereas this proliferative capacity is largely lost in adults across most species. The limited ability of adult cardiomyocytes to re-enter the cell cycle is a major cause of cardiac injury-induced morbidity and mortality. Here, we report that post-prandial Burmese python cardiomyocytes activate cell cycle re-entry to promote persistent cardiac growth. Burmese pythons normally eat large meals infrequently, resulting in reversible cardiac hypertrophy. We found that frequent feeding of large meals amplifies the modest post-prandial cardiac proliferation identified in an infrequent feeding interval. By activating E2F and Forkhead Box M1 (FoxM1) pro-proliferation transcriptional networks, frequently fed Burmese pythons initiate cardiomyocyte hyperplasia. These findings identify hyperplasia as a natural means of sustained cardiac growth in Burmese pythons and support the use of pythons as a new model for investigating proliferative cardiac remodeling.

## Introduction

Adult mammalian cardiomyocytes achieve cardiac growth via hypertrophy. This cellular growth augments cardiac function and can be enhanced by exercise and other stimuli through activation of physiological growth signaling pathways in cardiomyocytes (*1*). Meanwhile, pathological insults such as myocardial infarction (MI) cause irreversible cardiac tissue damage, maladaptive cardiac remodeling, and dysfunction, as adult mammalian cardiomyocytes have a limited ability to proliferate and replace damaged cells (*1*). The post-prandial Burmese python heart represents a rapid and robust model of physiological remodeling. Burmese pythons evolved as ambush-hunting predators and can both consume meals greater than their body mass and survive through months-long fasts during food scarcity. The python heart increases in mass by up to 40% within three days following consumption of large prey, a magnitude and rate of hypertrophy development that are unparalleled in other species (*2*, *3*).

Our earlier studies demonstrated that post-prandial cardiomyocyte hypertrophy in Burmese pythons is driven by activation of mammalian target of rapamycin (mTOR) and phosphatidylinositol 3-kinase (PI3K)-AKT signaling (*3*), while activation of Forkhead Box O1 (FoxO1) regulates the subsequent regression of hypertrophy (*4*). Although most of the post-prandial cardiac growth in Burmese pythons is reversible, the heart weight does not fully regress back to its pre-feeding size, which suggests the basal cardiac mass grows incrementally with each meal (*3*, *5*). In the current study, we detected modest cardiomyocyte hyperplasia in post-prandial Burmese python hearts during late digestion, a response that is potentiated by frequent feeding and contributes to long-term cardiac growth. We also demonstrated the proliferative effect could be translated to mammalian cardiomyocytes via circulating factor(s) in post-prandial plasma. These findings demonstrate that Burmese python cardiomyocytes engage in both hypertrophic (early digestion) and hyperplastic (late digestion) signaling post-prandially.

## Results

### Hyperplastic growth in adult Burmese python hearts

Previously, we concluded that the cardiac growth in post-prandial Burmese pythons was achieved through cellular hypertrophy instead of hyperplasia since we did not observe a proliferative signal (BrdU) at 1-day post-feeding (DPF) whereas we observed proliferation in the intestine (*3*). This is consistent with the observation in mammals that cardiomyocyte proliferation only occurs during embryonic and early neonatal stages. After birth, mammalian cardiomyocytes quickly become terminally differentiated and largely lose their proliferative capacity to re-enter the cell cycle (*6*, *7*). This cardiac maturation process involves a metabolic switch from glucose to fatty acids and hypertrophic growth (*7*). Only a few vertebrate species, including newts, zebrafish, and the recently identified spiny mouse, are known to reactivate cardiac mitotic signaling and cardiomyocyte proliferation as adults in response to cardiac injuries (*8–10*).

Our recent study demonstrated increased cell cycle gene enrichment in 4DPF Burmese python hearts (*4*), which implies the proliferative potential of some cardiac cell type. To verify whether mitosis was activated in cardiomyocytes, we co-immunostained python ventricular tissue sections with an antibody against phospho-histone H3 (pHH3), a late-stage cell cycle marker that is activated in mitosis, and an antibody against α-actinin (ACTN2), a sarcomere protein specifically expressed in cardiomyocytes. We found that the enhanced transcription of cell cycle factors previously observed at 4DPF preceded an over 10-fold increase in proliferative activity in 6DPF python hearts (Fig. 1A, 1B).

**Fig. 1.**
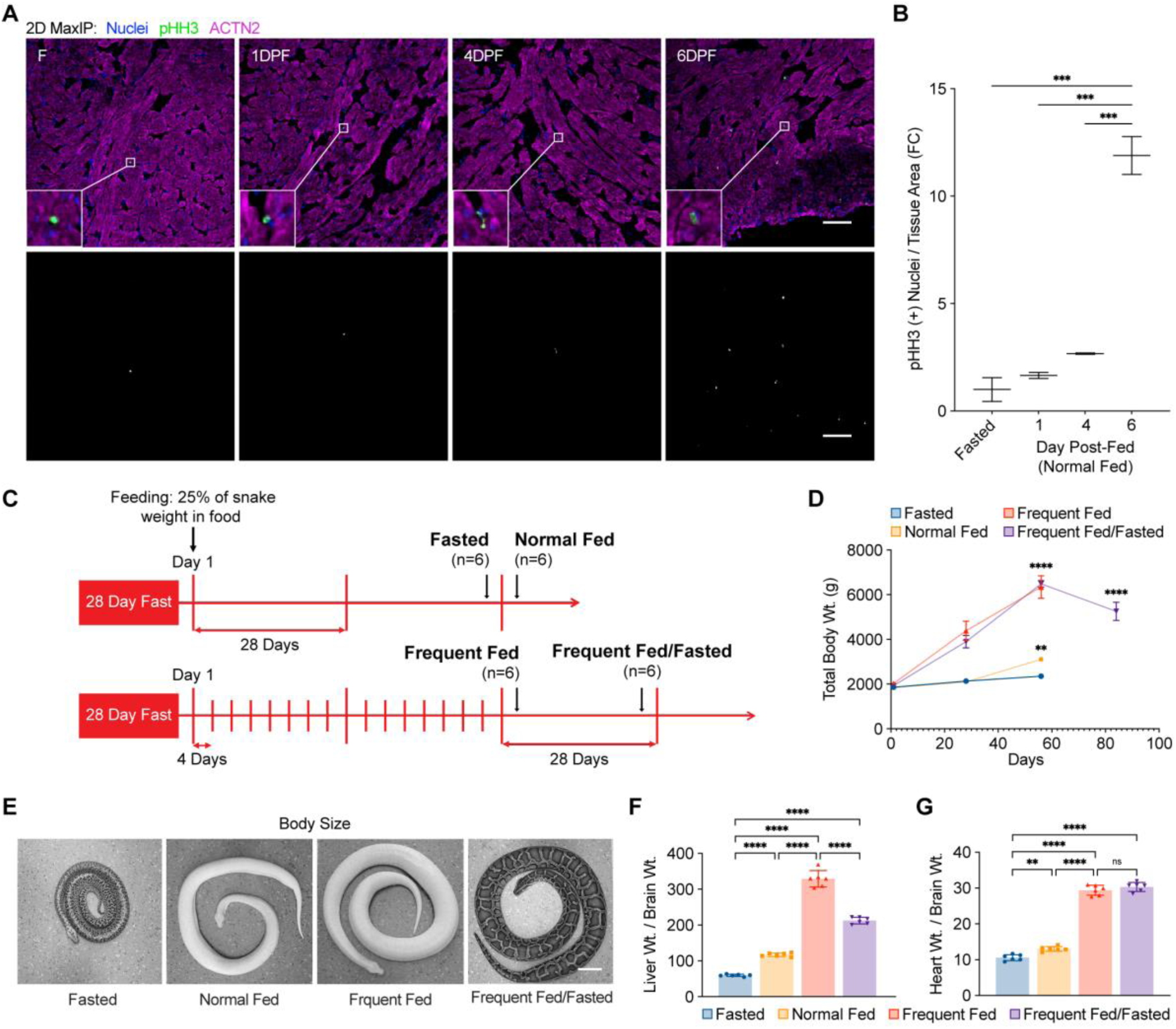
Frequent fed Burmese pythons experience sustained cardiac growth. **(A)** Immunohistochemistry staining of phospho-histone H3 (pHH3, green) and alpha actinin 2 (ACTN2, magenta) in python ventricle sections. Scale bar, 100 µm. The lower panel is the corresponding gray scale image for the pHH3 channel, day post-fed (DPF). **(B)** Quantification of pHH3(+) nuclei normalized to tissue area and plotted as a fold-change (FC). Data are represented mean ± SD, n=2. **(C)** Schematic showing feeding regimen for Burmese pythons: Fasted, Normal Fed, Frequent Fed, and Frequent Fed/Fasted. **(D)** Total body weight (Wt.) in the four groups over the course of the experiment. **(E)** Representative images of pythons in each group at the end of the study; scale bar, 10 cm. **(F-G)** Liver weight (D) and heart weight (E) normalized to brain weight (Supplemental Fig. 1). Data are represented mean ± SD, n=6 animals per condition. Ordinary one-way ANOVA with Tukey’s post-hoc test for multiple independent pairwise comparisons to the F group if it is not indicated; *p ≤ 0.05; **p ≤ 0.01; ***p ≤ 0.001; ****p ≤ 0.0001.

With increased meal frequency, we hypothesized that Burmese pythons would experience a sustained cardiac growth stimulus and an amplified proliferative response. Therefore, we designed a frequent feeding regimen. This approach highlights both a greater feeding frequency and a greater amount of food consumption. Burmese pythons were randomly assigned to either a ‘Fasted (four-week fast)’, or ‘Normal Fed (one meal after a four-week fast)’, ‘Frequent Fed (fed every four days for eight weeks)’ or ‘Frequent Fed/Fasted (fed every four days for eight weeks and then fasted for four weeks)’, with all meal sizes equivalent to 25% of python body mass (Fig. 1C). Throughout the feeding regimen, all groups showed steady growth after each meal.

After 16 consecutive feedings spanning two months, Frequent Fed pythons nearly tripled their body weight and cardiac mass compared to the Normal Fed pythons (Fig. 1D, 1E). With subsequent fasting in Frequent Fed/Fasted pythons, body weight and major organ masses, including the liver and kidney, significantly regressed. However, the cardiac ventricle and total heart weights remained elevated after fasting for one month (Fig. 1F, 1G, and fig. S1A, S1B). The fact that the heart did not regress after one month of fasting, as occurs with normal feeding frequency (*3*, *5*), suggests that Frequent Fed python hearts activated differential cellular signaling and likely facilitated hyperplastic cardiac growth in addition to normal feeding-induced transient hypertrophy.

### Frequent feeding amplifies cardiomyocyte cell cycle re-entry

To investigate how frequent feeding differentiated cardiac growth from normal feeding, we performed bulk RNA-sequencing (RNA-seq) on mRNA from python ventricular tissue. Principal-component analysis (PCA) indicated a sharp difference between the fed (Normal Fed & Frequent Fed) and the fasted (Fasted & Frequent Fed/Fasted) groups, while there was strong similarity between Normal Fed and Frequent Fed and between Fasted and Frequent Fed/Fasted (Fig. 3A). We therefore first performed a combined analysis of the fed groups (Normal Fed & Frequent Fed) compared to the combined fasted groups (Fasted & Frequent Fed/Fasted) to determine the general feeding-specific transcriptional response. Differential gene expression analysis revealed 3019 upregulated genes and 2779 downregulated genes in response to feeding (fig. S2A). Gene Ontology Biological Process (GO-BP) analysis revealed that upregulated pathways in the combined fed groups were enriched in metabolic processes, such as ATP synthesis and oxidative phosphorylation. The downregulated pathways were centered around cytoplasmic translation and ribosome biogenesis (fig. S2B). The RNA ribosomal regulation of transcriptional response was trending down in fed pythons, implying a turning point for transitioning digestion into the next stage after cardiac growth peaked at 72h post-feeding (*3*, *4*).

Although the hearts from fed pythons shared transcriptional features of metabolic pathway regulation, there was also a pronounced difference between Frequent Fed and Normal Fed (Fig. 2A). Differential gene expression analysis revealed only ∼50% of genes in Frequent Fed python hearts overlap with Normal Fed python hearts when individually compared to the Fasted pythons (fig. S2C). To determine how Frequent Fed python hearts are molecularly distinct from Normal Fed python hearts, we performed differential gene expression analysis in Frequent Fed versus Normal Fed ventricles followed by GO-BP analysis. Like the Fed vs Fasted comparison, Frequent Fed python hearts showed downregulation of cytoplasmic translation and ribosome assembly, demonstrating an exaggerated dampening of the translational response (Fig. 3B). Strikingly, the top upregulated pathways in the Frequent Fed python heart were associated with cell-cycle signaling, including chromosome separation and mitotic nuclear division. (Fig. 2B). Across all feeding conditions, the gene sets associated with mitotic pathways were exclusively enriched in the Frequent Fed python hearts (fig. S3A, S3B).

**Fig. 2.**
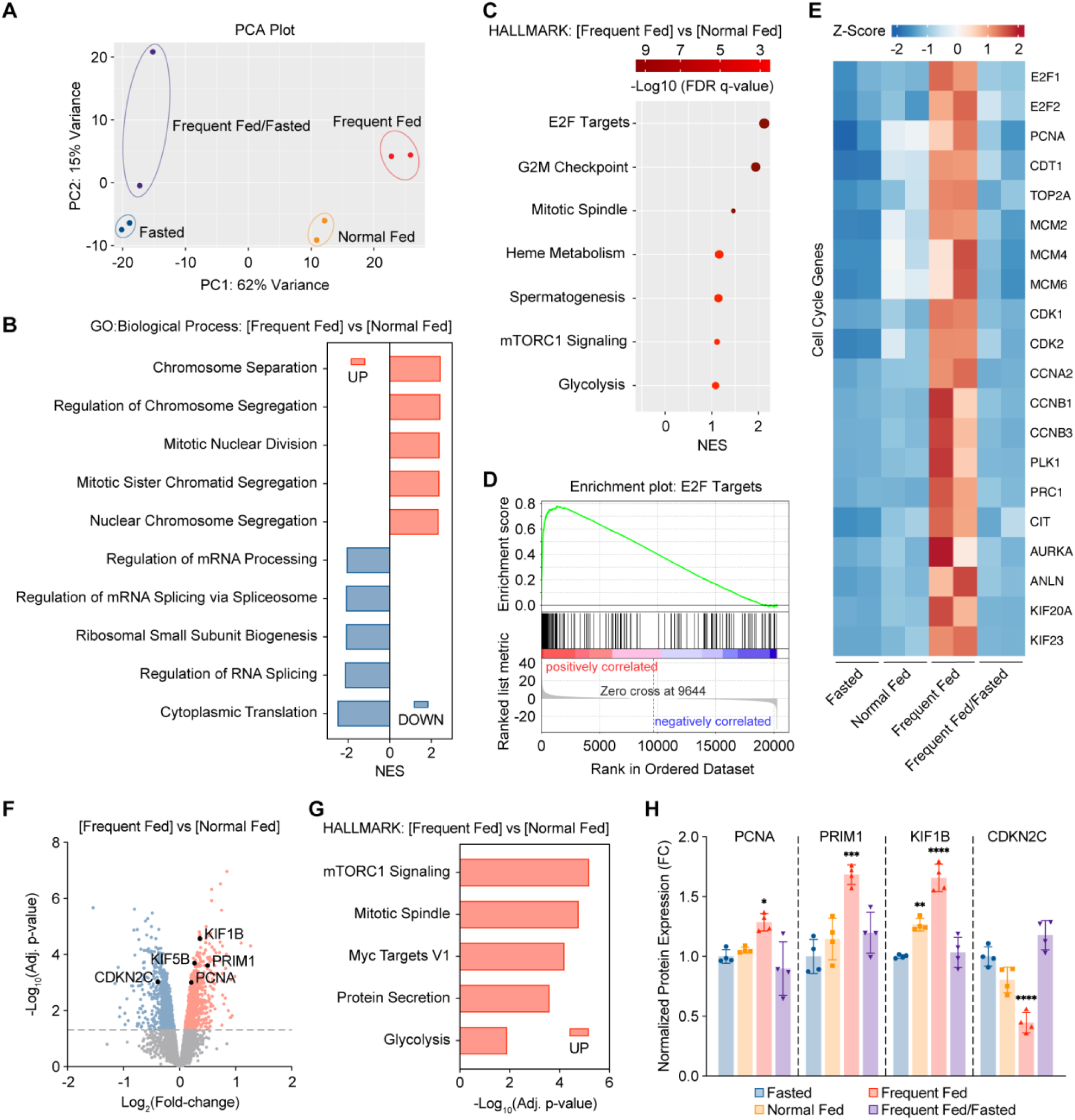
The frequent fed Burmese python cardiac transcriptome and proteome are enriched with cell cycle re-entry activity. **(A)** Principal component analysis (PCA) of the RNA-seq data from each feeding condition; red arrows denote comparisons of key groups discussed. n=2 animals per condition. **(B)** Gene Ontology (GO) Biological Process analysis of the up-regulated (UP) and down-regulated (DOWN) genes in Frequent Fed versus Normal Fed pairwise transcriptome comparison. Normalized enrichment score (NES). **(C)** Top enriched and up-regulated Hallmark pathways in Frequent Fed versus Normal Fed pairwise transcriptome comparison. **(D)** Gene set enrichment plot of E2F targets differentially expressed between Frequent Fed and Normal Fed pythons. **(E)** Analysis of the cell cycle genes differentially expressed across all feeding conditions highlighting exclusive enrichment in Frequent Fed pythons. **(F)** Proteomic analysis of protein expression in Frequent Fed versus Normal Fed pairwise comparison. **(G)** Top enriched and up-regulated Hallmark pathways in Frequent Fed versus Normal Fed pairwise proteome comparison. **(H)** Proteomic analysis of cell cycle protein expressions.

A recent study revealed that E2F transcriptional targets are distinctly enriched in embryonic mouse cardiomyocytes and mark their competency for entering the cell cycle (*12*). Applying the same gene set enrichment Hallmark pathway analysis to our differentially expressed genes, E2F Targets emerged as the top enriched pathway in Frequent Fed python hearts compared the Normal Fed python hearts (Fig. 2C, 2D). In addition, the enrichment of mTOR Signaling and Glycolysis suggested Frequent Fed python hearts have more robust growth signaling and reprogram to fetal-like metabolism (i.e., using glucose as a substrate) to support proliferative cardiac growth. Notably, 48 of the top 50 differentially expressed genes in the E2F Targets pathway were exclusively enriched in the Frequent Fed python hearts (fig. S4). Similarly, cell cycle genes including E2F targets (CNA, CDT1, TOP2A, MCMs), cell cycle G2/M regulators (CDK1, CDK2, CCNA2, CCNB1/3, PLK1), and cytokinesis markers (PRC1, CIT, AURKA, ANLN, KIFs) were exclusively upregulated in the OF python hearts (Fig. 2E and fig. S5). Additionally, expression of epithelial cell transforming 2 (ECT2), an essential gene for initiating cytokinesis (*13*), was significantly increased in the Frequent Fed python hearts, and the GO-BP analysis of Frequent Fed versus Normal Fed exhibited significant enrichment in Mitotic Cytokinesis (fig. S6A, S6B). Accordingly, our transcriptomic analysis strongly supports that adult Burmese python cardiac cells enter mitosis, and the cell cycle re-entry signatures are amplified by frequent large meals.

To verify transcriptional changes in cell cycle re-entry at protein level, we performed tandem-mass-tag (TMT) relative quantitative proteomics on Burmese python ventricular samples from each condition. We detected 322 significantly upregulated and 296 downregulated proteins in Frequent Fed versus Normal Fed pairwise comparison, including augmented expression of DNA replication markers (PCNA and PRIM1), cytokinetic kinesins (KIF1B and KIF5B), and decreased cell cycle inhibitor (CDKN2C) (Fig. 2F, 2H). Similar to the RNA-seq Hallmark pathway analysis in Frequent Fed versus Normal Fed, we identified increased expression of proteins involved in mTORC1 signaling, Mitotic spindle, and Myc target V1, indicating robust protein synthesis and cell cycle activity in the Frequent Fed python hearts (Fig. 2G). In contrast to downregulated cytoplasmic translation and RNA processing in transcriptomic analysis described above, our proteomics analysis revealed enhanced protein translation machinery in the Frequent Fed animals (fig. S7). The lag between transcription and translation suggests there are layers of molecular regulation in python hearts. Despite the lag between transcription and translation, these findings demonstrate aligned evidence at both transcriptomic and translational levels in supporting increased mitotic activity in Frequent Fed python cardiomyocytes.

### Dynamic cardiomyocyte growth by hypertrophy and hyperplasia

The absence of regression in heart size in Frequent Fed/Fasted pythons suggests long-term cardiac muscle mass accumulation, which aligns with there being an increase in total cardiomyocyte number. The expression of pro-proliferation transcription factors including E2F, TOP2A, MCMs, FoxM1 and other S phase cell cycle markers, including PCNA and MKI-67, were upregulated in both Normal Fed and Frequent Fed python hearts (fig. S5A-S5E, S9). Transcriptomic clustering analysis in Normal Fed versus Fasted also revealed a cluster (Cluster 1) of upregulated genes highly enriched in S phase genes of DNA replication, and their expressions were strongly enhanced in the Frequent Fed group (fig. S8). Along with the mitotic and cytokinetic gene expression and protein enrichment exclusive to Frequent Fed python hearts, these data provide compelling evidence to support accelerated and amplified mitotic cardiac growth in Frequent Fed pythons. However, the bulk analyses of the transcriptome and proteome include multiple cell types, so we turned to high resolution microscopy experiments to specifically examine the proliferative potential of python cardiomyocytes.

Consistent with the RNA-seq mitotic pathway enrichment, the co-immunostaining of pHH3 and ACTN2 showed a ∼40-fold increase in pHH3 signal in Frequent Fed python cardiomyocytes comparing to the Fasted python (Fig. 3A, 3B). In addition to hallmarks of cardiomyocyte proliferation, the Frequent Fed group showed significant cardiomyocyte hypertrophy as indicated by the decreased number of nuclei per tissue area (Fig. 3B), while cell size returned to baseline in the Frequent Fed/Fasted group despite the sustained cardiac growth observed (Fig. 1E, 3B). Through 3D imaging, we captured numerous dividing adult Burmese python cardiomyocytes at high resolution. The representative image showed a pair of pHH3(+) nuclei embedded in the cardiac muscle tissue was dividing and separating apart from each other (Fig. 3C, 3D, Movie S1). Although the mitotic marker pHH3 has its drawbacks as it does not fully distinguish cytokinesis from karyokinesis, the finding that cardiac mass remained elevated in the Frequent Fed/Fasted group, while cardiomyocyte hypertrophy was restored to baseline (Fig. 3B), indicates that the sustained cardiac growth induced by frequent feeding must be a result of increased cardiomyocyte number. Taken together, these data indicate that frequent large meal consumption directed python cardiomyocytes into a dynamic new phase: they not only undergo hypertrophy, but also enhance the existing post-prandial proliferative response and achieve a higher degree of cardiac growth to meet the increasing metabolic demand of their rapidly growing bodies. This further verifies that normal feeding activates cardiomyocyte proliferation, and that hyperplasia is a natural means of cardiac growth in adult python hearts. Notably, this proliferation occurs during late stages of digestion (6DPF) after the peak of hypertrophy has begun to regress (Fig. 1A, 1B).

**Fig. 3.**
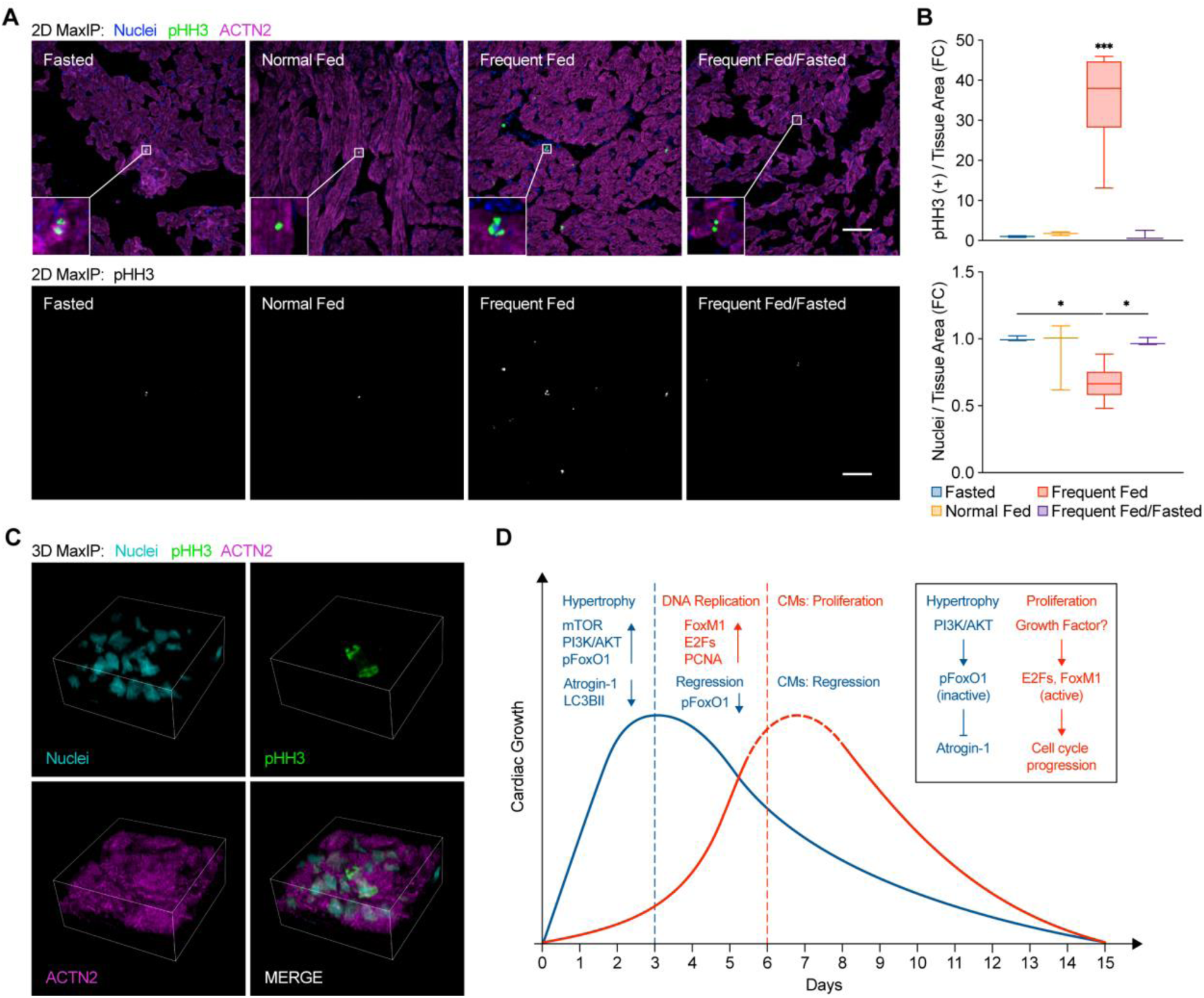
Frequent feeding accelerates proliferative cardiac growth in Burmese python hearts. **(A)** Immunohistochemistry staining of phospho-histone H3 (pHH3, green) and alpha actinin 2 (ACTN2, magenta) in python ventricle sections. Scale bar, 100 µm. The lower panel is the corresponding gray scale image for the pHH3 channel. **(B)** Quantification of pHH3(+) nuclei and total number of nuclei normalized to tissue area and plotted as a fold-change (FC). Data are represented mean ± SD, n=3 (Fasted, Normal Fed and Frequent Fed/Fasted), n=6 (Frequent Fed); ordinary one-way ANOVA test; *p ≤ 0.05; ***p ≤ 0.001. **(C)** Higher magnification image of a dividing cardiomyocyte identified in python ventricular tissue presented in 3D maximum intensity projection (MaxIP). Scale bar, 10 µm. **(D)** Model for post-prandial cardiac growth in Burmese python cardiomyocytes. Data are represented mean ± SD, n=6 animals per condition. Ordinary one-way ANOVA test with Tukey’s post-hoc test for multiple independent pairwise comparisons to the F group if it is not indicated; *p ≤ 0.05; **p ≤ 0.01; ***p ≤ 0.001; ****p ≤ 0.0001.

Given these findings, the current model of post-prandial hypertrophic growth of the Burmese python heart should be revised to a new dynamic model involving both cardiomyocyte hypertrophy and hyperplasia. First, the Burmese python heart exhibits rapid cardiac hypertrophy to accommodate the sudden rise in cardiac demand following a meal (1-3DPF), and this process is achieved via activation of mTOR and PI3K/AKT mediated growth (*3*) and suppression of Atrogin-1/FBXO32 mediated atrophy (fig. S9). While cardiac hypertrophy peaks at 3DPF in the Normal Fed paradigm, python cardiomyocytes initiate E2F and FoxM1 pro-proliferation transcriptional networks to promote DNA replication in later phases of digestion (4-6DPF). During the regression of post-prandial cardiac hypertrophy, a small population of cardiomyocytes enter M phase of cell cycle to proliferate (Fig. 1A, 1B, 3D). We expect this burst of proliferation following a meal contributes to lasting basal cardiac mass that steadily increases as the python grows. However, the length of this proliferative time window remains to be determined (indicated by dashed line section of the red curve in Fig. 3D).

### Frequent fed python plasma promotes hypertrophy and cell-cycle re-entry in mammalian cardiomyocytes

Previously, we demonstrated post-prandial Burmese python plasma promoted physiological cardiac hypertrophy in mammalian cardiomyocytes (*3*).To explore whether the cell cycle re-entry observed in the fed python heart was also caused by circulating factors, we took a similar approach and found 10% post-prandial python plasma treatment not only significantly promoted cellular hypertrophy, indicated by decreased nuclei count per cell area, but also activated cell cycle re-entry at 72 hours in neonatal rat cardiomyocytes comparing to the 10% fasted plasma, 10% calf serum, and serum free conditions (Fig. 4A, 4B, 4C). Normal fed plasma induced a >8-fold increase in cell cycle re-entry, which was further enhanced with frequent fed plasma (>12-fold increase) (Fig. 4B). We also captured numerous mammalian cardiomyocytes in different phases of mitosis indicated by cytokinesis marker Aurora kinase B (AURKB) (Fig. 4D). More importantly, the robust cardiac cell cycle re-entry response is specific to post-prandial python plasma as fasted plasma and calf serum largely induced cellular hypertrophy only. Calf serum is commonly supplemented in the media to promote cell growth. Despite observing abundant pHH3(+) signals from the calf serum condition, most of these signals were found in non-cardiomyocytes. This demonstrates the cell cycle re-entry observed in the post-prandial python heart can be translated to mammalian cardiomyocytes, likely via cardiomyocyte-specific circulating factor(s) in the plasma.

**Fig. 4.**
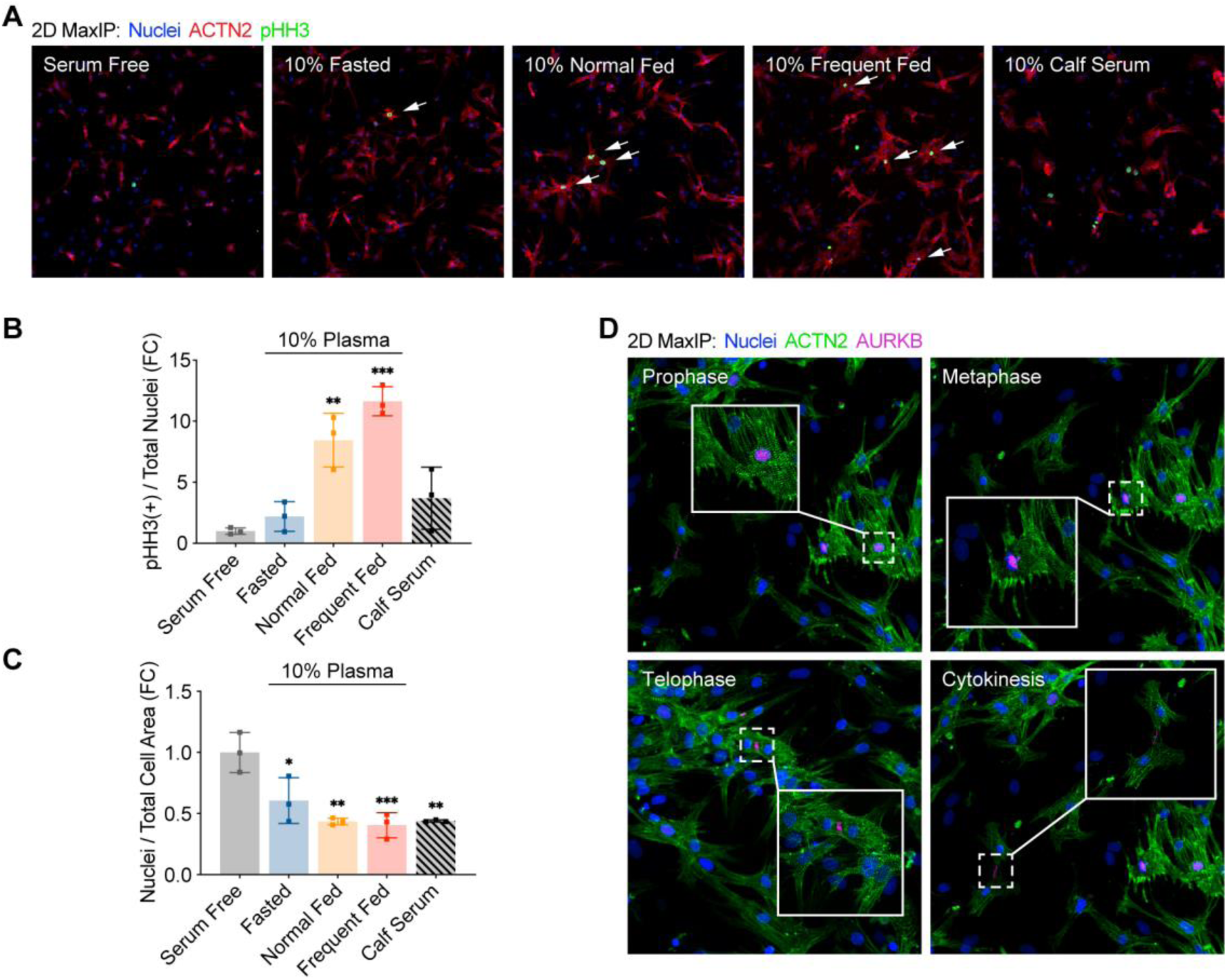
Post-prandial python plasma promotes hypertrophy and cell-cycle re-entry in mammalian cardiomyocyte. **(A)** Immunofluorescent staining of phospho-histone H3 (pHH3, green) and alpha actinin 2 (ACTN2, red) in neonatal rat cardiomyocytes. (**B-C**) Quantification of pHH3(+) nuclei and total number of nuclei normalized to cell area and plotted as a fold-change (FC). **(D)** Immunofluorescent staining of aurora kinase B (AURKB, magenta) and alpha actinin 2 (ACTN2, green) captured neonatal rat cardiomyocytes in different mitotic phases after treated by frequent fed python plasma. Data are represented mean ± SD, n=3 per condition. Ordinary one-way ANOVA test with Tukey’s post-hoc test for multiple independent pairwise comparisons to the Serum Free group if it is not indicated; *p ≤ 0.05; **p ≤ 0.01; ***p ≤ 0.001; ****p ≤ 0.0001.

## Discussion

In this study, we introduced a novel frequent feeding paradigm in Burmese pythons to test the proliferative potential of their cardiomyocytes. Knowing that almost all adult mammalian cardiomyocytes achieve postnatal cardiac growth via hypertrophy rather than hyperplasia, our results led to the discovery of striking distinctions between Burmese pythons and mammalian hearts: adult Burmese python cardiomyocytes activate hyperplastic cardiac growth during meal digestion, and the proliferative activity is significantly potentiated by increasing meal frequency. Our data, therefore, add Burmese pythons to the short list of vertebrate species known to have cardiomyocytes with proliferative potential, which includes newts, zebrafish and spiny mice (*8–10*). Like newts and zebrafish, Burmese pythons are also ectotherms and exhibit low baseline metabolic rates (*14*)). However, unlike these previously studied species, Burmese python hearts exhibit hyperplasia as natural means of cardiac growth rather than injury-induced cardiac regeneration and repair. Compared to the cardiac dedifferentiation process during post-injury cardiac regeneration in zebrafish (*15*, *16*), we observed similar sarcomere disassembly, upregulation of fetal genes (ACTA1, TNNT1) and downregulation of genes involved in sarcomere maintenance (TCAP) (fig. S10), whereas there is a lack of injury-induced cardiac regeneration signaling in the Burmese python heart compared to the cardiac apex resection in zebrafish. This suggests Burmese python cardiomyocytes evolved to utilize a different type of molecular regulation to achieve proliferation.

Moreover, we found post-prandial cardiac proliferation in pythons is marked by increased FoxM1 expression (fig. S9), which is akin to post-injury zebrafish cardiomyocytes and proliferative neonatal rat cardiomyocytes (15). FoxM1, a proliferation-associated transcription factor, is essential for zebrafish cardiomyocytes to complete regeneration after injury (*17*). Additionally, cardiomyocyte-specific loss of FoxM1 leads to cell-cycle withdrawal and decreased neonatal rat cardiomyocyte proliferation (*18*). During this process, FoxM1 has been shown to play an antagonistic regulatory role with AKT/FoxO1 signaling during cardiac growth (*18*). Similar to mammalian cardiomyocytes, we recently reported that the activity of FoxO1 plays a central regulatory role in controlling post-prandial cardiac hypertrophy via Atrogin-1/FBXO32 and autophagy-mediated proteolysis in Burmese python hearts (*4*). Here, we propose a similar molecular regulation: immediately after meal ingestion, python cardiomyocytes activate PI3K/AKT signaling, which deactivates FoxO1 and downstream Atrogin-1 to promote robust cardiac hypertrophy. When cardiac demand starts to decrease, FoxO1 becomes re-activated and drives cardiac regression (*4*). Meanwhile, increased E2F and FoxM1 target gene expression promotes cell cycle progression and proliferation, whereas the upstream proliferative growth signaling remains to be identified (Fig. 3D). On the other hand, cellular proliferation by dedifferentiating existing cardiomyocytes leads to sarcomere disassembly and therefore reduced contractile function. Notably, overexpression of cytokinesis regulators (PLK1 and ECT2) in adult mammalian hearts resulted in widespread postnatal cardiomyocytes mitosis, but also contractile dysfunction (*19*).Frequent feeding significantly induced PLK1 and ECT2 expression in python hearts and resulted in 0.22% cell cycle re-entry rate at this fixed time, which translates to a projected rate of 2.63% per year (Fig. 2E, Table S1). In a normal feeding cycle, cell cycle progression initiates when cardiac demand decreases (6DPF). This separates the need for basal cardiac growth (hyperplasia at 6DPF) at the expense of reduced contractile function from the high cardiac demand during early digestion (hypertrophy only at 1-3DPF). During frequent feeding, we expect the sustained, robust cardiac hypertrophy can functionally compensate for the amplified cell cycle re-entry rate.

Collectively, this study extends our molecular understanding of the dynamic post-prandial cardiac remodeling in Burmese pythons. Our data support that hyperplastic growth during late stages of digestion contributes to long-term increases in cardiac mass. The low rate of post-prandial hyperplasia continues during adulthood to steadily increase basal cardiac mass as the python grows from centimeters in length as a juvenile to upwards of 6 meters in adulthood. Compared to existing models of cardiac regeneration (e.g., zebrafish), Burmese pythons are more complex vertebrates with stronger cardiac function, and their hearts are unique in that they maintain low pulmonary blood pressure while produce high systemic blood pressure akin to mammals (*20, 21*). They have two atria and a partially septated ventricle, giving them a more structurally similar cardiac anatomy to mammals (*20, 21*). Lastly, we demonstrated translational potential of circulating factor(s) in the post-prandial python plasma to promote mammalian cardiomyocyte proliferation. Frequent fed python plasma potently induced cell cycle re-entry in neonatal rat cardiomyocytes; however, the specific circulating factor(s) and its origin remain to be determined. Taken together, our findings elevate the Burmese python as a novel platform to investigate molecular mechanisms that regulate adult cardiomyocyte cell cycle re-entry.

## Materials and Methods

### Animals and tissue collection

Burmese pythons (n=24) were obtained as hatchlings from Bob Clark Reptiles (Oklahoma City, OK) and housed individually at ∼28°C on a 12/12-hour light/dark cycle. Pythons were fed a meal equivalent to 25% of their body mass on a biweekly basis until they reached 1300-1500g. Four months prior to the beginning of the study, each python was placed on a 4-week (28 days) feeding regime. Each meal was 25% of the python’s body mass. After four months, pythons were randomly assigned to either a ‘Fasted’, or ‘Normal Fed’, or ‘Frequent Fed’ or ‘Frequent Fed/Fasted’ group with n=6. After the four-month acclimation period, both groups of the Fasted and Normal Fed pythons were maintained on a 28-day feeding regime for the remainder of the study (8 weeks). The Frequent Fed and Frequent Fed/Fasted groups were fed every 4 days over the same 8-week period. and the Frequent Fed/Fasted group was fasted for another 4 weeks following 8-week frequent feeding. Pythons were euthanized at the end of each prandial state: Normal Fed and Frequent Fed, 3 days after their final meal; Fasted and Frequent Fed/Fasted, 28 days after their final meal. Tissues and plasma were collected, snap-frozen, and stored at −80°C. All procedures were conducted under the approval of the University of Colorado, Boulder IACUC.

### Neonatal rat ventricular myocytes (NRVM) isolation and culture

Cardiomyocytes were isolated from one-day-old Sprague Dawley rat pups as previously described (*4*). Purified ventricular cardiomyocytes were plated at 250,000 cells per well on 12-well plates with glass coverslips. Approximately 24 hours after isolation, the wells were washed twice and then incubated in experimental media (MEM, 20 mM HEPES, 2 µg/mL vitamin B12) for 18 hours prior to the start of the experiment. NRVMs were treated with serum free experimental media, 10% Fasted python plasma, 10% Normal Fed python plasma, 10% Frequent Fed python plasma, and 10% calf serum with experimental media respectively at 0 hour and 24 hours for a total of 72 hours treatment.

### Immunohistochemistry

Python hearts were collected and embedded in O.C.T. Compound (Fisher Healthcare, 23730571), and cryosectioned at 10 μm intervals. For immunofluorescent staining, cryosections were air-dried for 15 min at room temperature and fixed with cold 4% PFA for 30 min. Section slides were then washed twice with PBS and permeabilized with 0.5% Triton X-100 (Fisher Scientific, BP151-500) in PBS for 20 min. Sections were then blocked in 2% BSA/0.25% Triton X-100/PBS blocking solution for 1 h at room temperature and stained with the indicated primary antibodies prepared in 2% BSA/0.125% Triton X-100/PBS at 4 °C overnight using the following dilutions: sarcomeric alpha-actinin (Thermo Fisher Scientific, MA1-22863, 1:250), pHH3 (Cell Signaling Technology, 9701, 1:200). Sections were subsequently washed with PBS three times (5 min for each wash), and incubated with corresponding secondary antibodies conjugated to Alexa Fluor 488 or 555 (Cell Signaling Technology) prepared in 2% BSA/0.125% Triton X-100/PBS at room temperature for 1.5 h. After secondary antibody incubation, sections were washed with PBS three times (5 min for each wash) and mounted using VECTASHIELD® PLUS Antifade Mounting Medium with DAPI (Vector Laboratory, H-2000).

### Immunocytochemistry

Wells with NRVMs were washed twice with PBS, and fixed with ice cold methanol for 1 min and then 4% PFA for 10 min. Fixed cells were washed twice with PBS and permeabilized with 0.5% Triton X-100 (Fisher Scientific, BP151-500) in PBS for 20 min followed by 0.1% Triton X-100 in PBS for another 30min. Permeabilized cells were then blocked in 0.5% BSA/0.1% Triton X-100/PBS blocking solution for 1 h at room temperature and stained with the indicated primary antibodies prepared in 0.5% BSA/0.1% Triton X-100/PBS at 4 °C overnight using the following dilutions: sarcomeric alpha-actinin (Thermo Fisher Scientific, MA1-22863, 1:250), pHH3 (Cell Signaling Technology, 9701, 1:200), and AURKB (Thermo Fisher Scientific, MA5-15321, 1:250). Cells were subsequently washed with PBS three times, and incubated with corresponding secondary antibodies conjugated to Alexa Fluor 488 or 555 (Cell Signaling Technology) prepared in 0.5% BSA/0.1% Triton X-100/PBS at room temperature for 1.5 h. After secondary antibody incubation, sections were washed with PBS three times and mounted using VECTASHIELD® PLUS Antifade Mounting Medium with DAPI (Vector Laboratory, H-2000).

### Microscopy and imaging analysis

Burmese python ventricles section: pHH3 stain, three field of views were randomly selected from each stained tissue slide. Images (10X 3x3) used for quantifications and representative images (10X 1x1) were obtained using a Nikon Spinning Disk confocal microscope. 3D images (100X 1x1) were obtained using a Nikon A1R Laser Scanning confocal microscope. Neonatal rat ventricular myocytes: pHH3 and AURKB stain, images (10X 3x3) used for quantifications, representative images (20X 1x1, 40X 1x1), and 3D images (100X 1x1) were obtained using a Nikon Spinning Disk confocal microscope. The data analysis and visualization work were performed at the BioFrontiers Institute’s Advanced Light Microscopy Core using the software pack Imaris (Oxford Instrument).

### RNA-sequencing and analysis

Total RNA was isolated from python tissue and cells with Trizol reagent. The concentration of RNA was determined using Qubit RNA BR Assay Kit (Invitrogen Q10211) and Qubit Flex Fluorometer. Frozen RNA samples were sent to Novogene for sequencing and analyzed based on Burmese python (*Python molurus bivittatus,* Taxonomy ID: 176946) genome. Deferential gene expression analysis was performed using R package DESeq2 (*22*). The gene set was then pre-ranked for Gene Ontology Biological Pathway analysis (c5.go.bp.v2023.1.Hs.symbols.gmt) using Gene Set Enrichment Analysis software with the Hallmark Pathway analysis (Molecular Signature Database, MSigDB) (*23, 24*). For those enriched pathway with 0 FDR-qval, 0 was assigned as 1.0*e-9 for graphing. RNA-seq analysis graphing profiles were generated using RStudio (v2023.12.1.402) and GraphPad Prism 10 (GraphPad Software Inc).

### Proteomics and analysis

The protocol for tandem-mass-tag (TMT) quantitative proteomics of python ventricular samples was previously described (cite PNAS). In brief, python ventricular tissue (n = 4 per group from Fasted, Normal Fed, Frequent Fed, and Frequent Fed/Fasted) was pulverized while frozen and then added to a 5% SDS/0.75% sodium deoxycholate 50 mM Tris (pH 8.5) buffer and immediately boiled for 10 minutes. The samples were then centrifuged at 12,000 x g for 10 minutes and the supernatant collected. Protein concentration was determined by BCA assay. Chris protocol. The protein samples were reduced, alkylated, and then digested using the SP3 method (cite). Tryptic peptides were then labeled with TMT-Pro 16-plex (Thermo Scientific) according to the manufacturer’s protocol. The peptides were then fraction with high pH reversed-phase C18 UPLC using a custom-packed UChrom column (1.8 µm 120Å). 12 fractions were eluted at 20 µL/minute using 2-50% acetonitrile using an M-class UPLC (Waters). Peptides were then lyophilized by vacuum centrifugation, resuspended in 3% (vol/vol) ACN, 0.1% (vol/vol) trifluoroacetic acid (TFA). LC-MS/MS analysis was performed using an Ultimate 3000 nanoUPLC (Thermo Scientific) and Q-Exactive HF-X mass spectrometer (Thermo Scientific) as previously described (4). The raw proteomics data files were searched against the *Python bivittatus* proteome database using MaxQuant (v.2.0.3.0) with cysteine carbamidomethylation as a fixed modification and methionine oxidation and N-terminal acetylation as variable modifications. Data were thresholded at 1% false discovery rate (FDR) and fold changes and p-values were calculated with limma using an R-script.

### Quantitative real time PCR (RT-qPCR) analysis

Total RNA was extracted from python cardiac muscle tissue with Trizol and reversed transcribed using Super Scrip III (Invitrogen, 12574018) with random primers. mRNA levels were quantified with SYBR Green (Invitrogen, 4312704). Due the dramatic changes in gene expression profile in post-prandial Burmese python hearts, all conventional normalizers such as 18s were changing. Therefore, we determined concentration of RNA using m ore precise chemical reaction-based method, Qubit RNA BR Assay Kit (Invitrogen, Q10211) and Qubit Flex Fluorometer rather than wavelength-based Nanodrop. Based on the precisely measured concentration of RNA, the same amount of cDNA (5 ng) was loaded for each RT-qPCR reaction, and these experiments were performed in triplicate. RT-qPCR primers were designed using whole genomic and transcriptomic Burmese python sequence on NCBI (Taxonomy ID: 176946). RT-qPCR primers are available in Table S1.

### Western blot

Burmese python ventricle was pulverized in liquid nitrogen cooled pulverizer, lysed overnight in RIPA buffer (Cell Signaling Technology, 9806S), supplemented with protease and phosphatase inhibitors cocktail (Thermo Scientific Technology, 8442), centrifuged at 15,000 *g* for 15 minutes at 4°C, and the supernatant was collected from each sample and stored at −80°C. The concentration of protein lysates was quantified using DC protein assay kit (Bio-Rad, 5000112). 25 μg of protein from the lysate was resolved by NuPAGE™ 4-12%, Bis-Tris gel (Invitrogen, WG1403A), transferred on to nitrocellulose membrane (Bio-Rad, 1620115), blocked in 5% fat-free milk/0.2% TWEEN-20/TBS at room temperature for 1h and stained with the indicated primary antibodies prepared in 5% BSA/0.2% TWEEN-20/TBS at 4°C overnight using the following dilutions: LC3B (Cell Signaling Technology, 2775), 1:1000. Membranes were subsequently washed with TBS three times (5 min for each wash), and incubated with corresponding HRP-linked secondary antibodies (Anti-rabbit IgG, Cell Signaling Technology, 7074) prepared in 5% BSA/0.2% TWEEN-20/TBS at room temperature for 1.5 h. After secondary antibody incubation, membranes were washed with TBS three times (5 min for each wash). Membranes were imaged after 1-minute incubation of Western Lightning Plus-ECL (PerkinElmer, NEL104001EA). Images were then analyzed using ImageQuant.

### Annual cell-cycle re-entry rate calculation

Cell cycle re-entry rate at the time of tissue collection was calculated by pHH3(3) nuclei per total nuclei. The annual rate is estimated by the following equation: annual rate = (1 + fix rate)^12^.

### Statistical analysis

Statistical analyses were performed using GraphPad Prism 10 (GraphPad Software Inc), with p-value < 0.05 considered significant unless otherwise indicated. All data were analyzed for outlier before running statistical tests. Detailed information on the statistical analysis is supplied in the figure legends. All data are displayed as means ± SDs; each data point indicates a biological replicate, unless otherwise indicated.

## Acknowledgments

We thank Dr. Massimo Buvoli and members of the Leinwand laboratory for helpful discussion. We thank Mary Allen for technical support on RNA-sequencing data analysis. We thank IACUC staff at BioFrontiers Institute for support with animal experiments. We thank Dr. Kristen Bjorkman, Dr. Claudia Crocini, and Dr. Massimo Buvoli for editing the English language of the manuscript.

## Author Contributions

Conceptualization: YT, AKP, RD, BH, LAL

Methodology: YT, TGM, AKP, CCE, RD, BH

Investigation: YT, TGM, AKP, CDO

Visualization: YT

Funding acquisition: TGM, LAL

Project administration: LAL

Supervision: LAL

Writing – original draft: YT

Writing – review & editing: YT, TGM, LAL

## Data and materials availability

The raw RNA sequencing data will be deposited in the NCBI Gene Expression Omnibus repository. The raw proteomics data will be deposited in the Proteome Xchange Consortium via the PRIDE partner repository.

## Figures

**Fig. S1.**
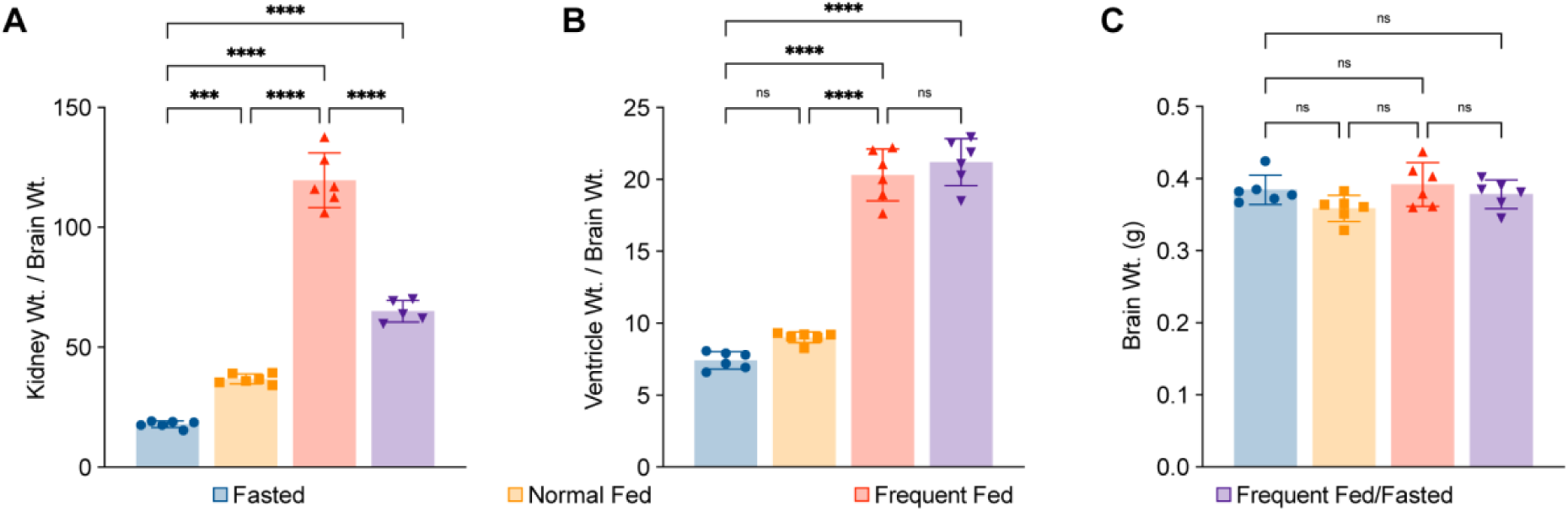
**(A)** Kidney weight normalized to brain weight. Brain weight. **(B)** Ventricle weight normalized to brain weight. **(C)** Brian weight. Data are represented mean ± SD, n=6 animals per condition. Ordinary one-way ANOVA with Tukey’s post-hoc test for multiple independent pairwise comparisons to the F group if it is not indicated; *p ≤ 0.05; **p ≤ 0.01; ***p ≤ 0.001; ****p ≤ 0.0001.

**Fig. S2.**
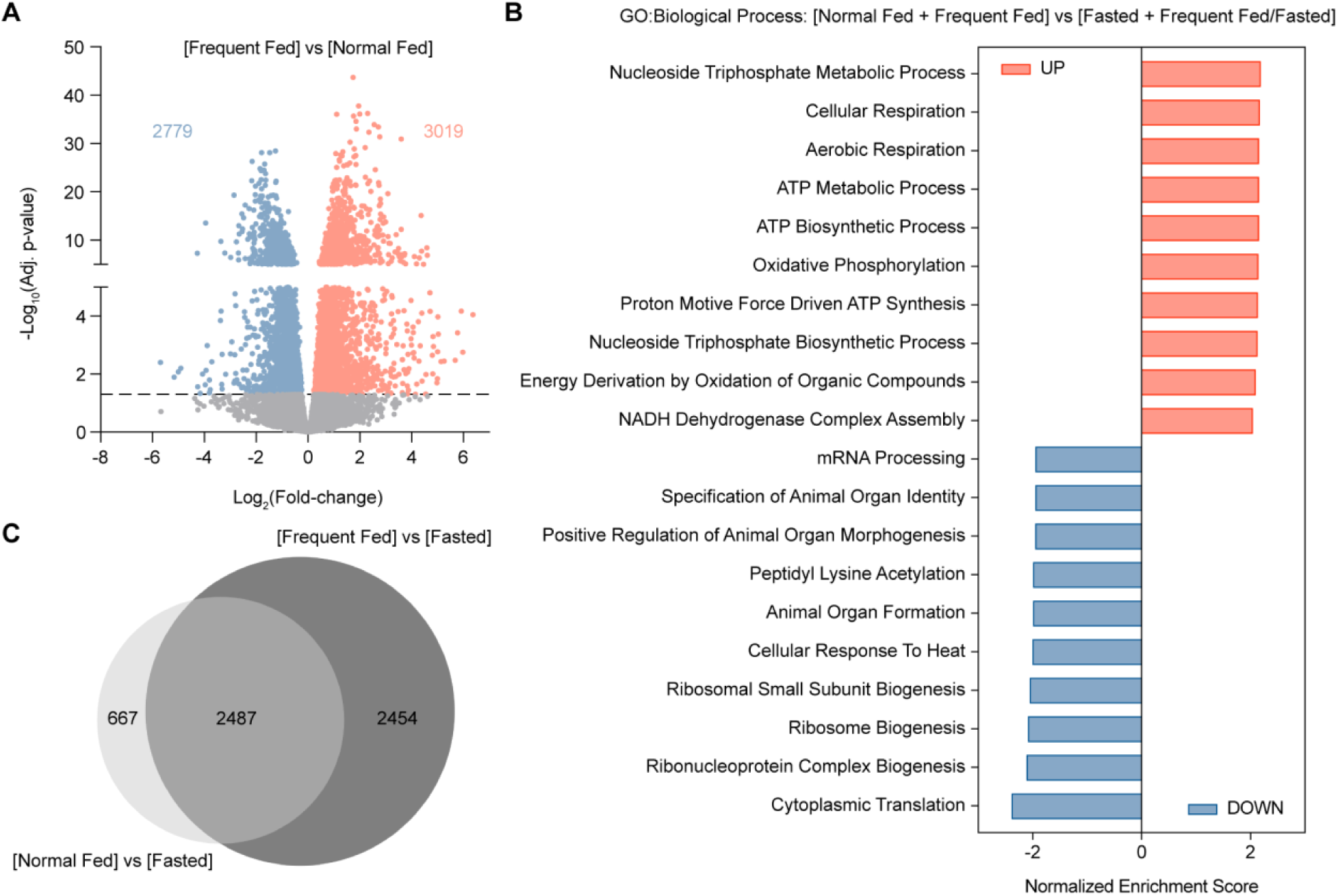
**(A)** Volcano plot of differentially expressed genes (DEG) in the fed groups [Normal Fed + Frequent Fed] compared with the fasted groups [Fasted + Frequent Fed/Fasted]. DEG cut-off: p-value ≤ 0.05. **(B)** Gene Ontology Biological Process (GO-BP) analysis of the up-regulated (UP) and down-regulated (DOWN) genes in [Normal Fed + Frequent Fed] versus [Fasted + Frequent Fed/Fasted] pairwise transcriptome comparison. **(C)** Pie chart of overlapping DEG between Frequent Fed and Normal Fed compared to the Fasted control.

**Fig. S3.**
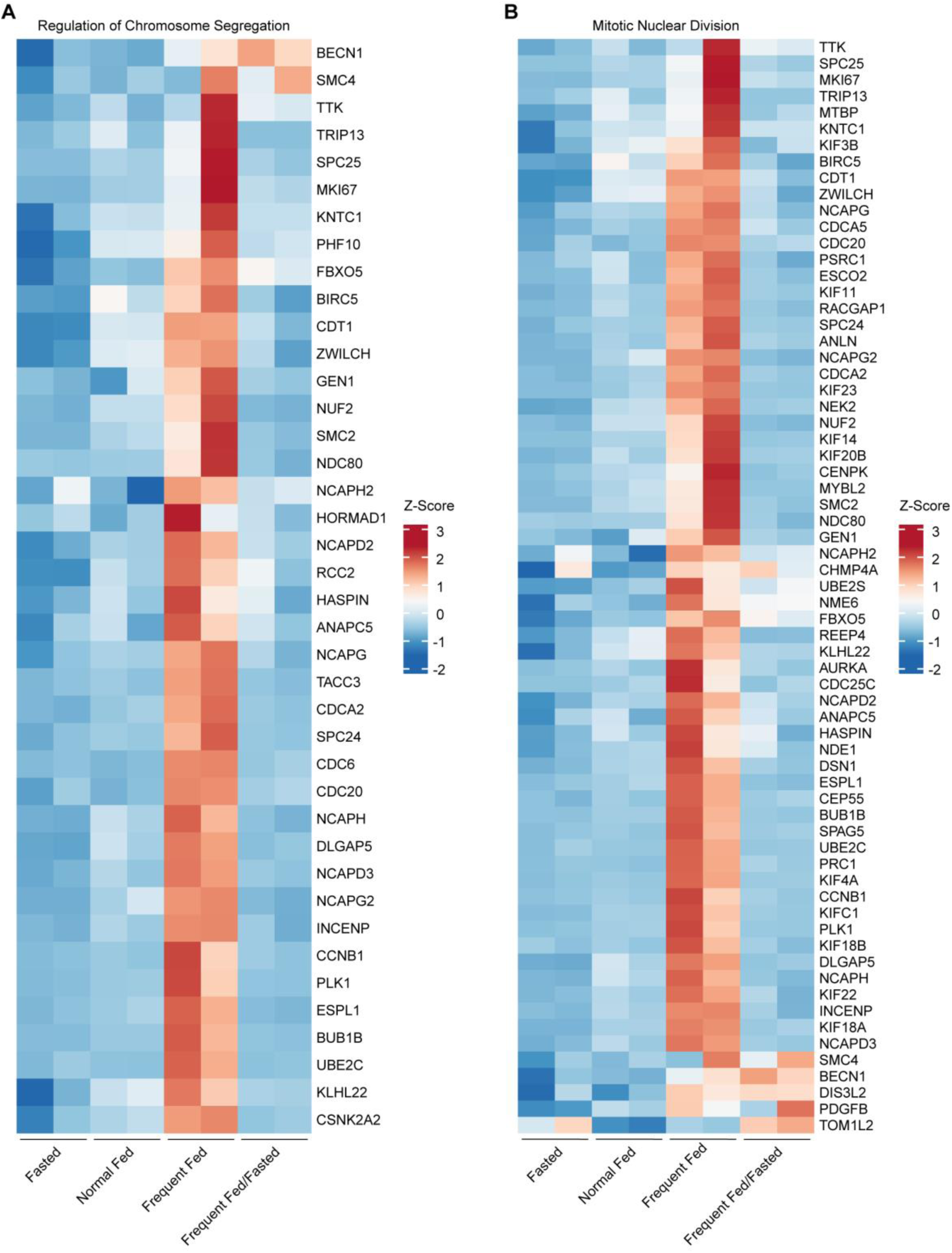
**(A)** Heatmap of top GO-BPs, Regulation of Chromosome Separation. **(B)** Heatmap of top GO-BPs, Mitotic Nuclear Division.

**Fig. S4.**
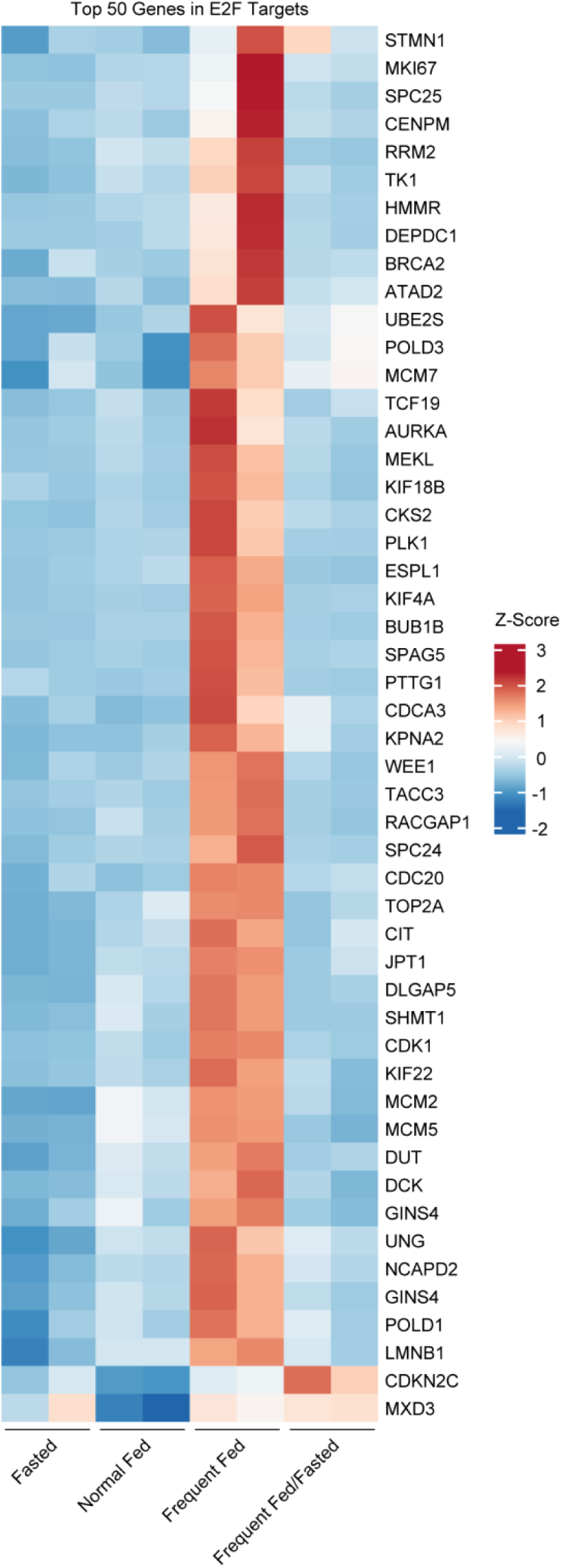
Analysis of the top 50 E2F targets DEGs across all feeding conditions highlighting exclusive enrichment in Frequent Fed pythons.

**Fig. S5.**
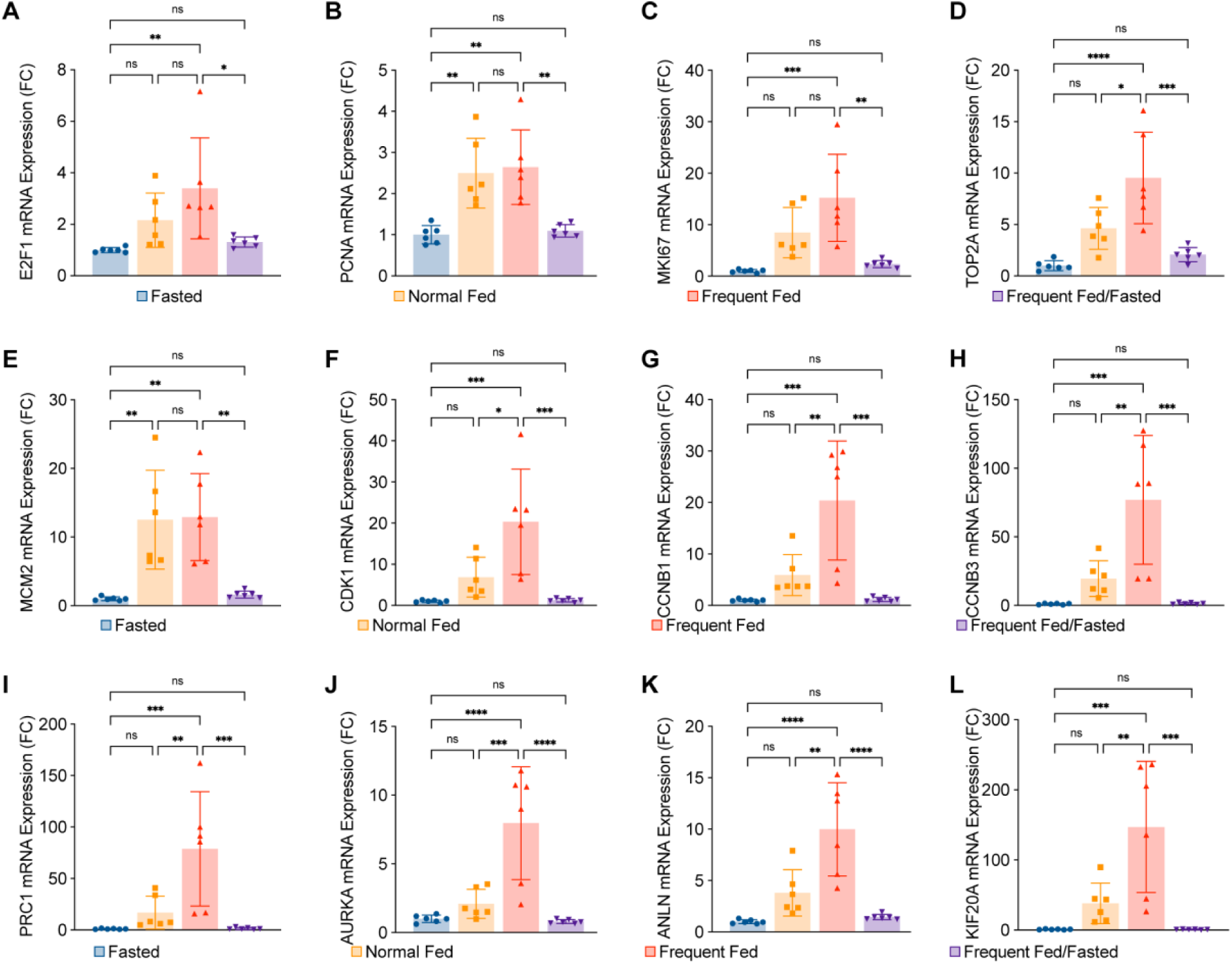
Quantitative PCR analysis for genes involved in cell cycle: **(A-E)** E2F targets. **(F-H)** G2/M. **(I-L)** Cytokinesis. Data are represented mean ± SD, n=6 animals per condition. Ordinary one-way ANOVA with Tukey’s post-hoc test for multiple independent pairwise comparisons to the F group if it is not indicated; *p ≤ 0.05; **p ≤ 0.01; ***p ≤ 0.001; ****p ≤ 0.0001.

**Fig. S6.**
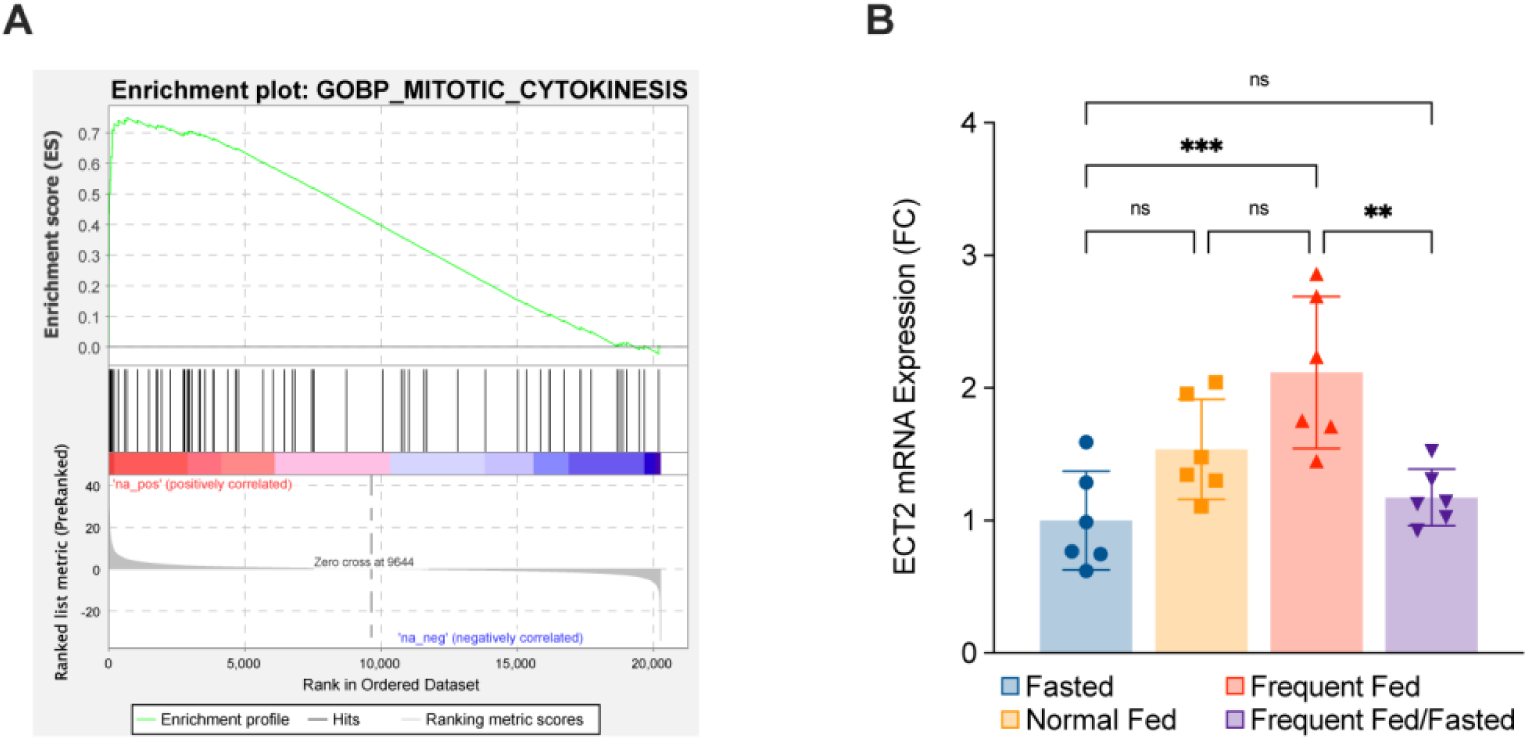
**(A)** Enrichment plot of Gene Ontology Biological Process (GO-BP), Mitotic Cytokinesis. **(B)** Quantitative PCR analysis for cytokinesis gene, ECT2. Ordinary one-way ANOVA with Tukey’s post-hoc test for multiple independent pairwise comparisons to the F group if it is not indicated; *p ≤ 0.05; **p ≤ 0.01; ***p ≤ 0.001; ****p ≤ 0.0001.

**Fig. S7.**
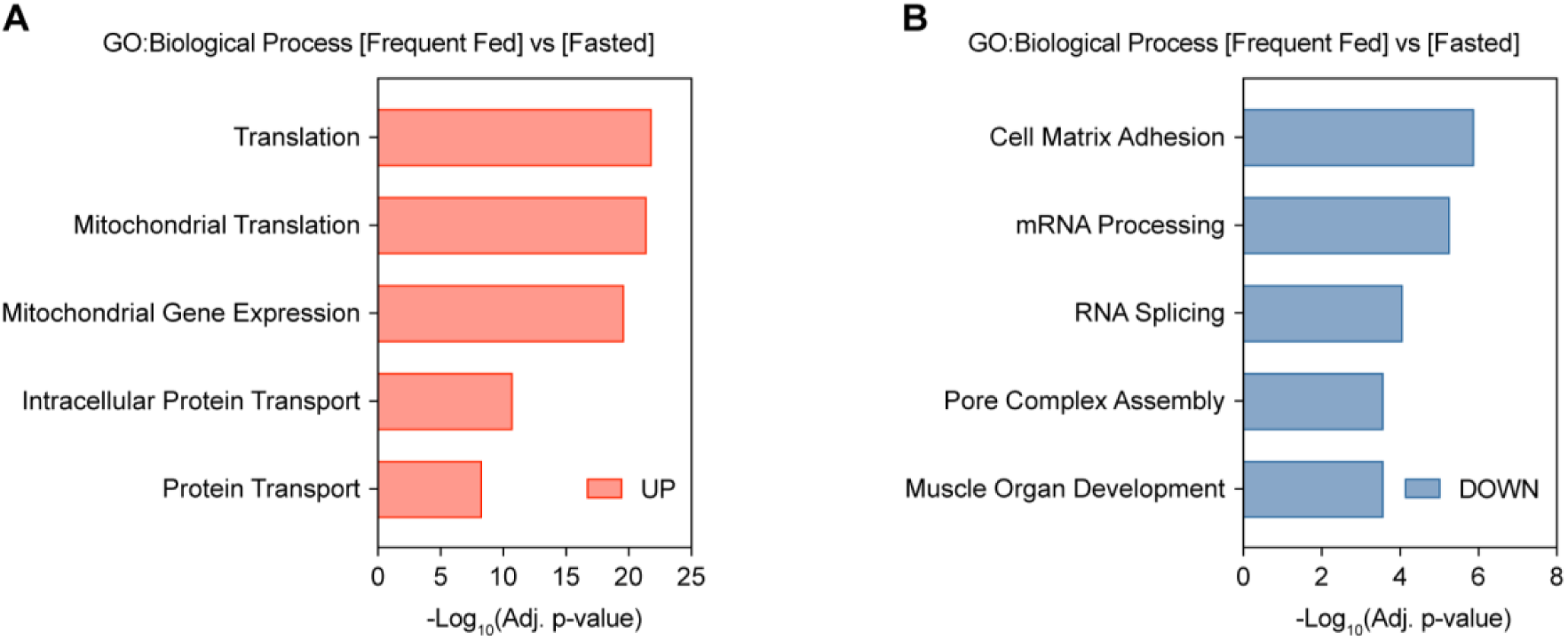
GO-BP analysis of the up-regulated (UP) proteins **(A)** and GO-BP analysis of the down-regulated (UP) proteins **(B)** in Frequent Fed versus Fasted pairwise comparison.

**Fig. S8.**
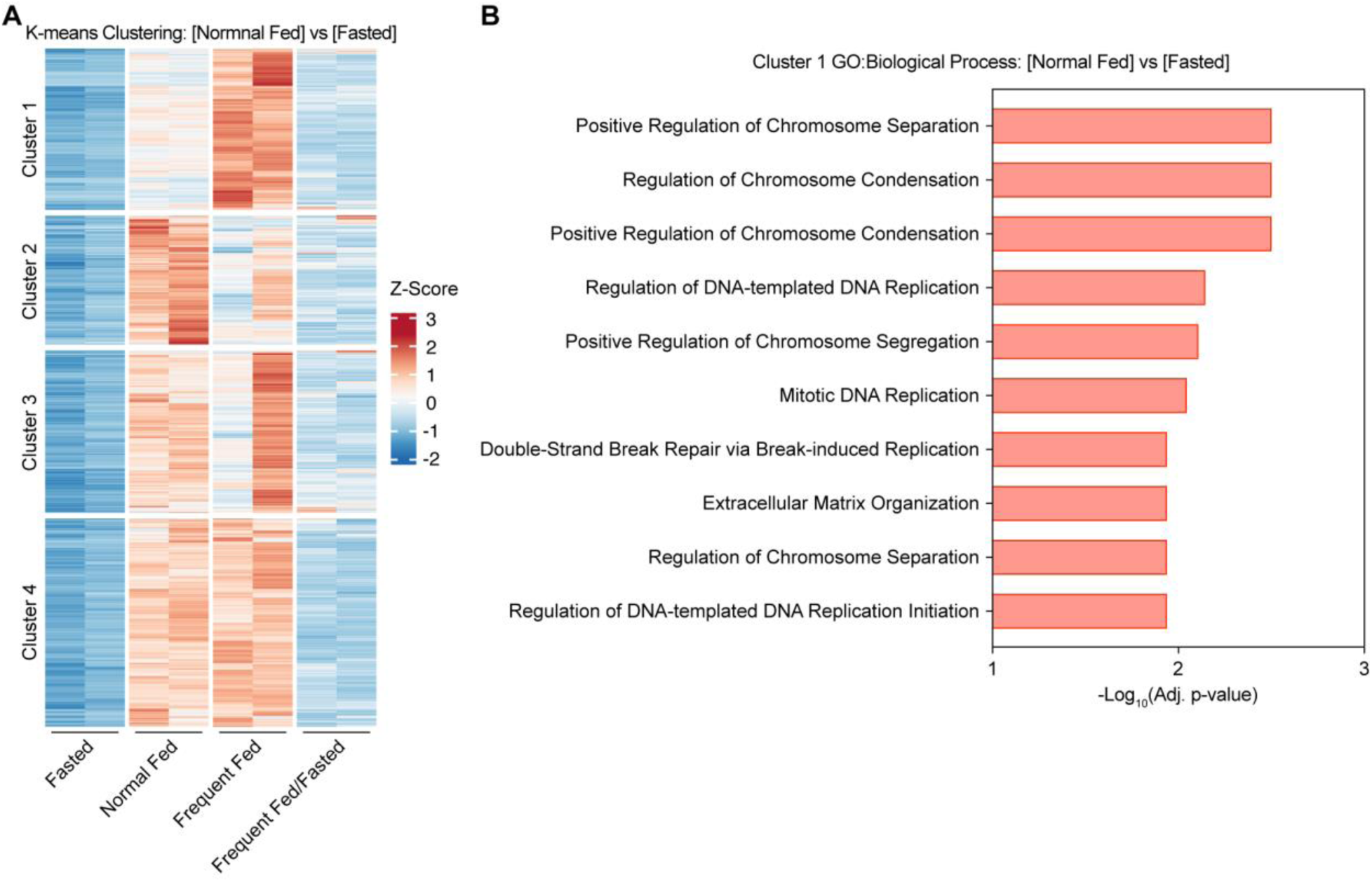
**(A)** K-means clustering analysis of upregulated DEGs in Normal Fed versus Fasted pairwise comparison. across all feeding conditions highlighting exclusive enrichment in Frequent Fed pythons. **(B)** GO-BP analysis of the up-regulated (UP) genes in Cluster 1 identified in (A).

**Fig. S9.**
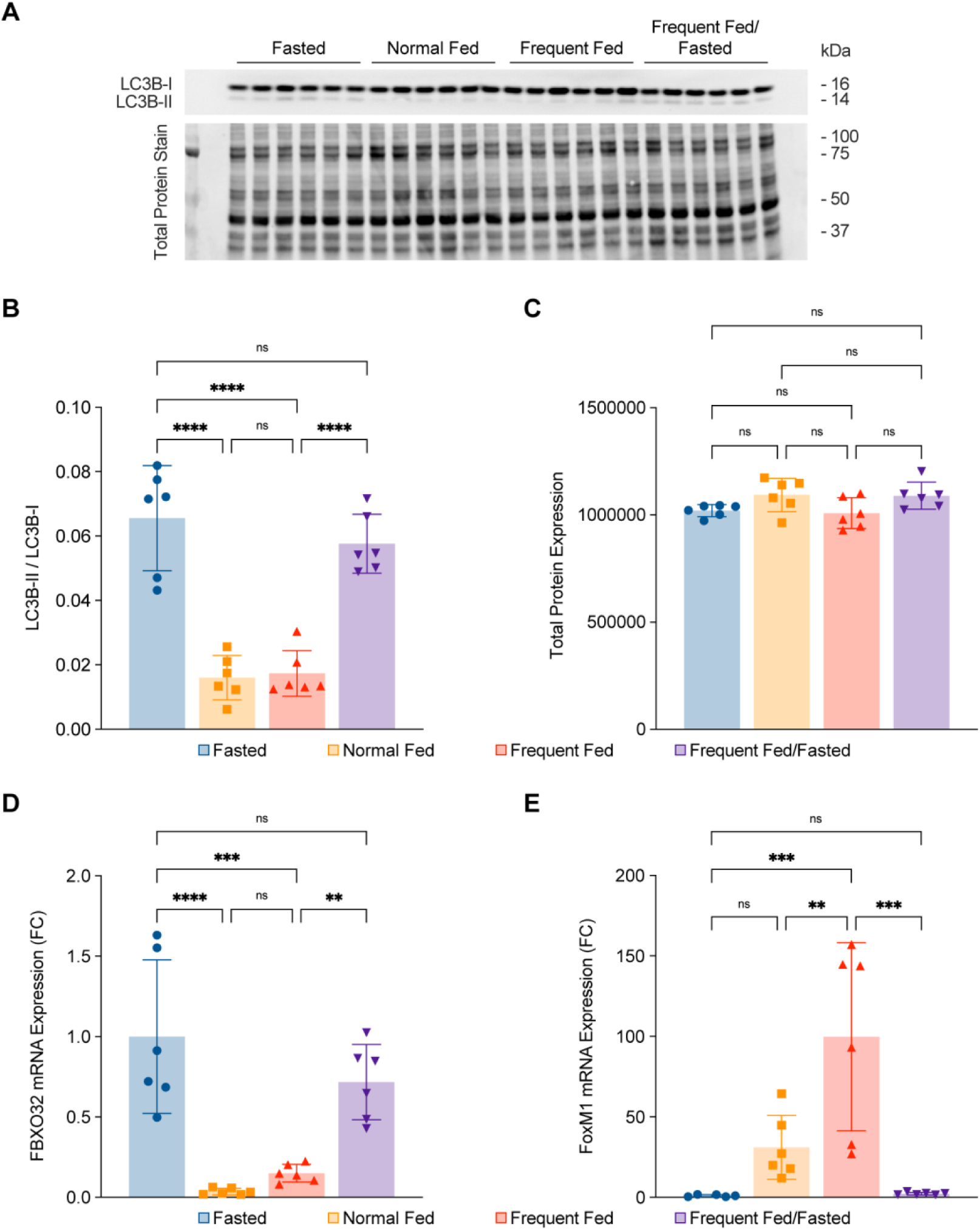
**(A)** Western blot for LC3B-I and LC3B-II. **(B)** Quantification of LC3B-II/LC3B-I cleavage ratio. **(C)** Quantification of total protein expression for normalization. **(D)** Quantitative PCR analysis for atrophic genes, FBXO32. **(E)** Quantitative PCR analysis for pro-proliferation transcription factor, FoxM1. Data are represented mean ± SD, n=6 animals per condition. Ordinary one-way ANOVA with Tukey’s post-hoc test for multiple independent pairwise comparisons to the F group if it is not indicated; *p ≤ 0.05; **p ≤ 0.01; ***p ≤ 0.001; ****p ≤ 0.0001.

**Fig. S10.**
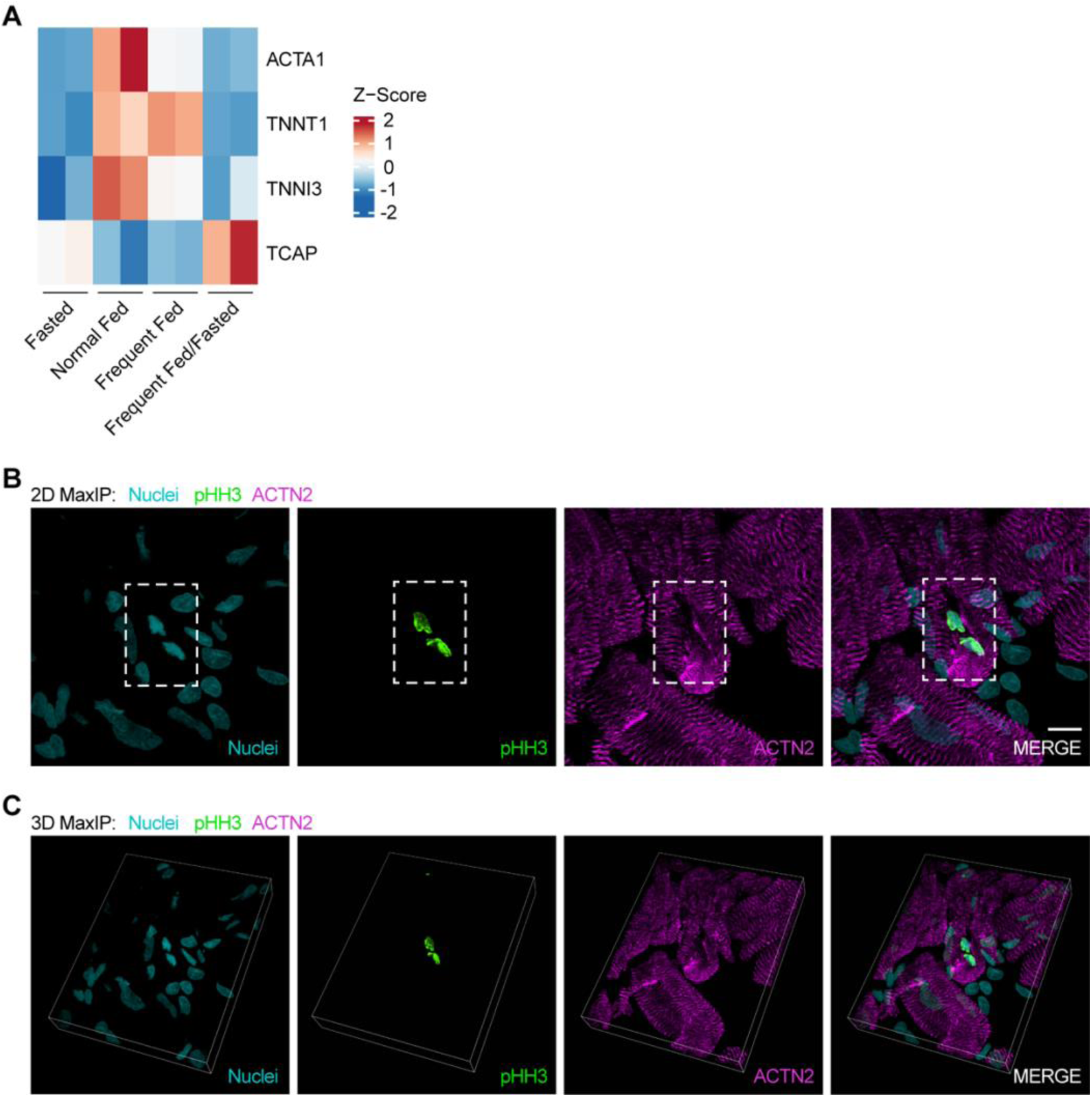
(**A**) Heatmap of sarcomere maintenance gene expression in RNA-seq transcriptomic analysis. (**B-C**) Sarcomere disassembly in a pHH3(3) Burmese python cardiomyocyte.

**Fig. S11.**
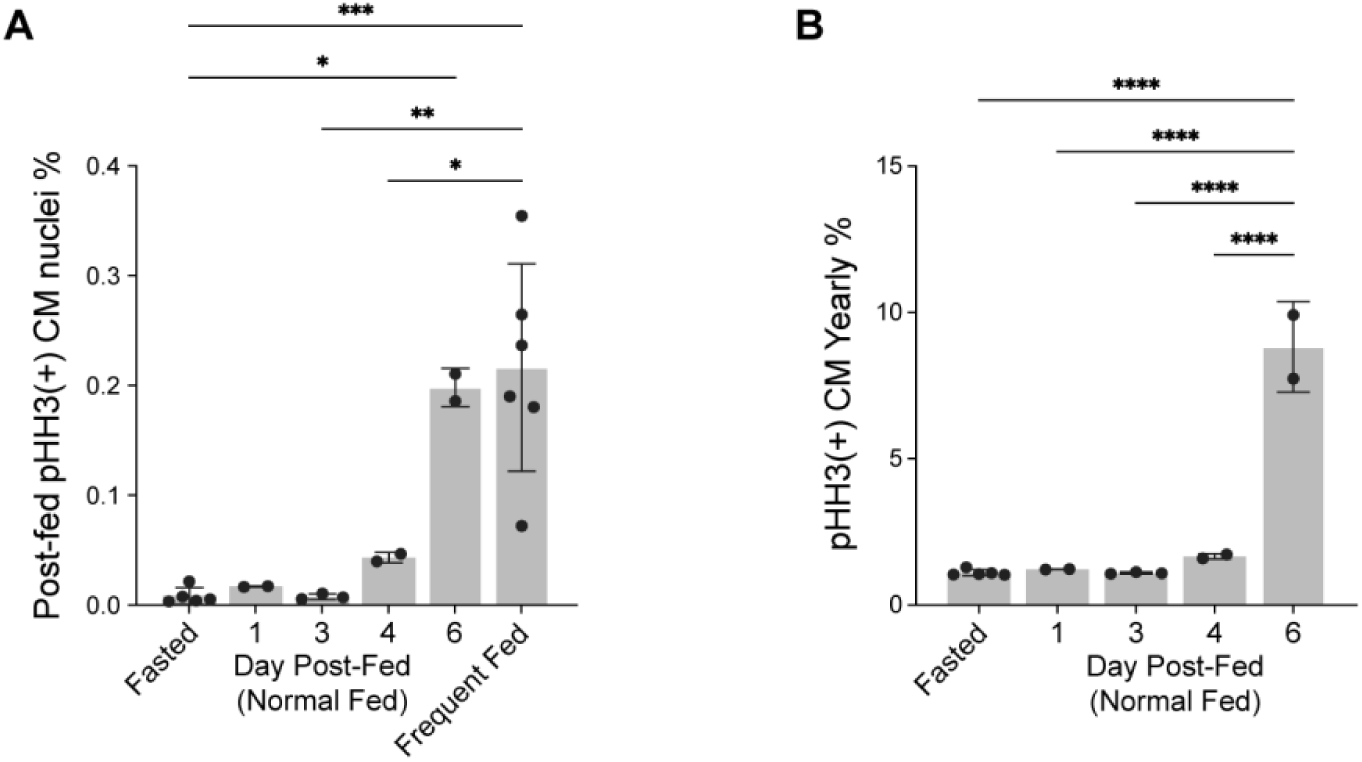
Burmese python cardiomyocytes cell cycle re-entry rate at the time of tissue collection (**A**) and projected annual rate (**B**).

**Table S1.** Cell cycle re-entry rate of Burmese python ventricular cardiomyocytes.

|  | **Fixed time point rate (%)** | **Projected yearly rate (%)** |
| --- | --- | --- |
| **Fasted (n=5)** | 0.0084 | 0.1006 |
| **1DPF (n=2)** | 0.0170 | 0. 2036 |
| **3DPF (n=3)** | 0.0076 | 0.0916 |
| **4DPF (n=2)** | 0.0433 | 0.5205 |
| **6DPF (n=2)** | 0.1983 | 2.4054 |
| **Frequent Fed (n=6)** | 0.2163 | 2.6321 |

**Table S2.**
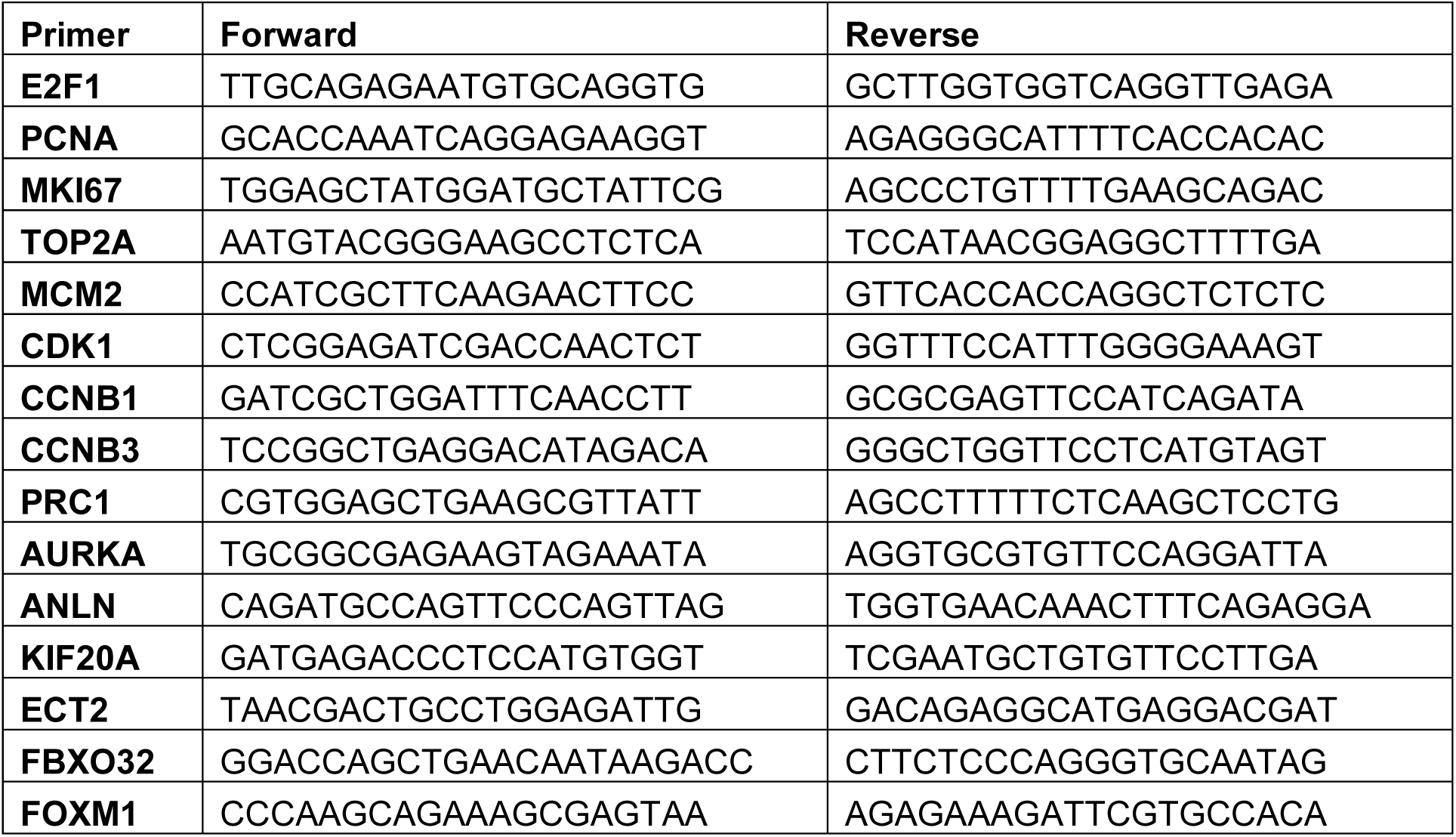
Quantitative PCR primer sequences.

## Movie S1

3D image construction of a dividing cardiomyocyte identified in python ventricular tissue presented in maximum intensity projection (MaxIP) at 100X.

